# Structural analysis reveals three families of bacterial tRNA mimics

**DOI:** 10.64898/2026.10.20.760346

**Authors:** Jia Wang, Pierre Boudry, Yuting Xie, Yuanlin He, Yangyi Ren, Qiqi Chen, Eric Westhof, David M J Lilley, Olga A. Soutourina, Lin Huang

## Abstract

Comparative genomics has identified over 1,000 candidate structured RNA motifs, yet most remain functionally uncharacterized. Here, we combine crystallography, comparative analysis, and bacterial genetics to identify three families of bacterial tRNA-like RNAs. The skipping-rope RNA adopts an L-shaped fold resembling tRNA despite distinct secondary-structure connectivity and tertiary interactions. In *Clostridioides difficile*, the skipping-rope RNA RCd1 is abundantly expressed and processed into a highly stable RNA species. RCd1 overexpression suppresses sporulation, whereas its deletion enhances early spore formation, supporting a role in sporulation control. Guided by the skipping-rope structure, we identify a conserved architectural core and use it to recognize additional candidate tRNA mimics. Crystal structures of DUF2693-FD and salivarius-1 RNAs confirm a shared tRNA-like scaffold assembled through distinct interaction networks. Together, these findings expand the repertoire of bacterial tRNA mimics and establish a structural framework for recognizing shared architectures among uncharacterized ncRNAs, linking one such family to bacterial developmental regulation.

## INTRODUCTION

Comparative genomics and transcriptome-wide analyses have uncovered hundreds of structured bacterial noncoding RNA (ncRNA) families, many of which remain functionally uncharacterized^1,2^. Conserved sequences and nucleotide covariation support their predicted secondary structures but often provide limited clues to their biological roles. Determining their three-dimensional structures can reveal relationships to characterized RNAs that are not apparent from sequence or secondary structure, providing a basis for classification and functional investigation.

Transfer RNA (tRNA) mimicry illustrates how distinct RNAs can share a recognizable three-dimensional architecture. Canonical tRNAs adopt an L-shaped fold stabilized by long-range interactions between the D and T loops, which form the characteristic tRNA elbow. This architecture also occurs in other RNAs: viral tRNA-like structures and bacterial transfer-messenger RNA (tmRNA) engage components of the translation machinery, whereas the bacterial yjdF riboswitch and Y RNA YrlA extend tRNA mimicry to regulatory and other noncanonical contexts^3–6^. These examples highlight the versatility of the tRNA scaffold and raise the question of which structural features are shared by RNAs with different sequences and secondary-structure topologies.

The skipping-rope motif is a widespread bacterial ncRNA family identified by comparative analysis of intergenic regions^1^. Its conserved helices and long-range pseudoknot suggest a defined tertiary fold, while the frequent presence of downstream intrinsic transcription terminators is consistent with expression as an independent RNA. One member, RCd1 from the human pathogen *Clostridioides difficile*, is an abundant intergenic transcript that associates with the RNA chaperone Hfq and has been implicated in sporulation control^7–9^. Despite these biological clues, the three-dimensional architecture of skipping-rope RNAs and the structural basis of their sequence conservation remain unknown.

To investigate the architecture underlying the conserved skipping-rope motif, we determined its crystal structure, which revealed an unexpected resemblance to tRNA. This finding prompted us to explore both the biological properties of a native family member and the occurrence of related architectures in other bacterial ncRNAs. Analysis of RCd1 connected the skipping-rope family to RNA processing, prolonged transcript stability, and negative regulation of sporulation. In parallel, structural features of skipping-rope RNA guided the identification and crystallographic characterization of two further families, DUF2693-FD and salivarius-1. Together, these structures show how different secondary-structure topologies and local interaction networks support a common tRNA-like core. The resulting framework extends the classification of these ncRNAs beyond sequence-based families and highlights structural relationships that would otherwise remain difficult to recognize.

## RESULTS

### The skipping-rope RNA adopts a tRNA-like L-shaped architecture

The skipping-rope RNA family (Rfam RF02924) was originally identified by comparative genomic analysis of bacterial intergenic regions^1,2^. Its consensus secondary structure comprises conserved helices and a covariation-supported long-range pseudoknot, with connectivity that does not readily resemble a known RNA fold^2^ (Figures 1A and S1A).

**Figure 1.**
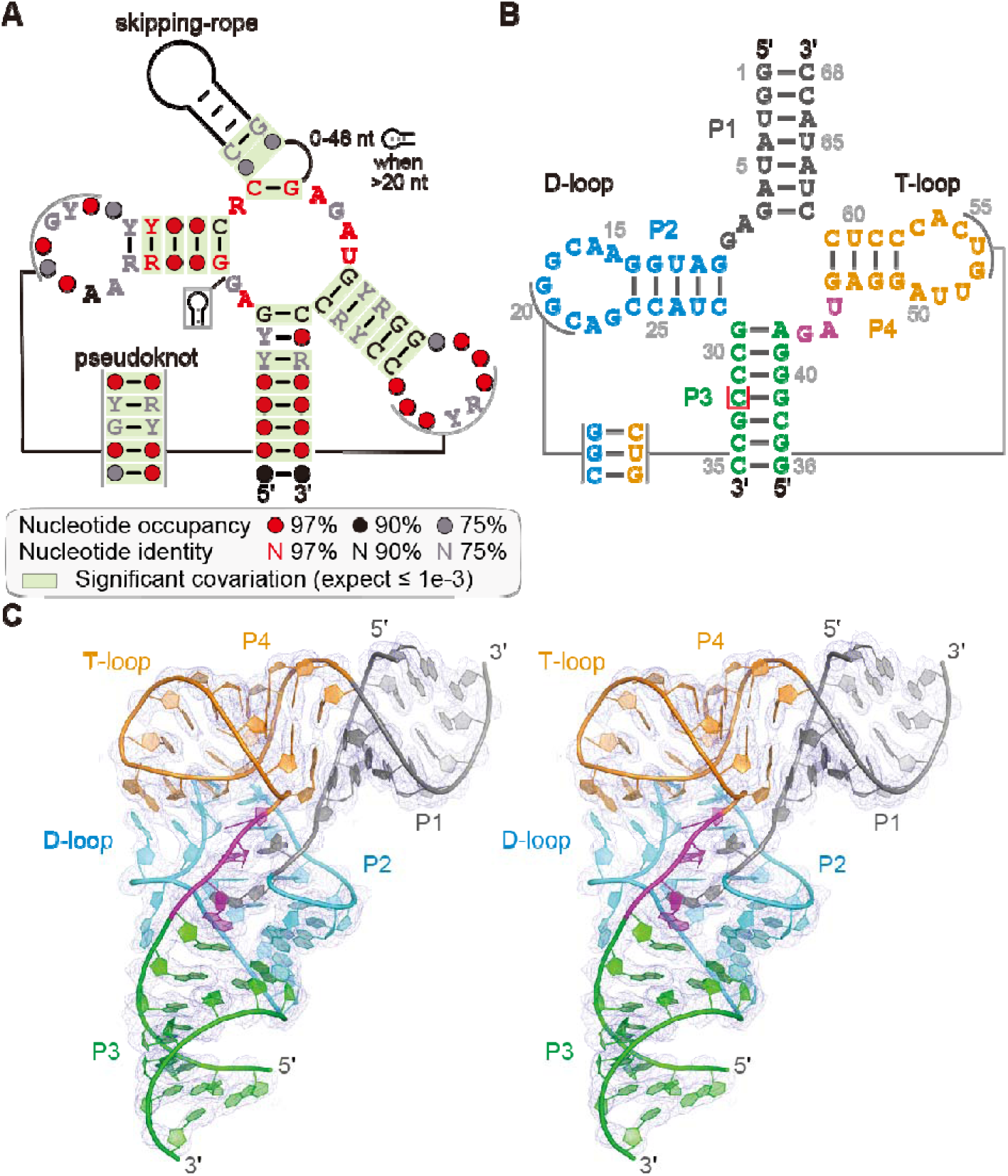
Crystal structure of the skipping-rope RNA reveals a tRNA-like L-shaped architecture. (**A**) Consensus sequence and secondary-structure model of the skipping-rope RNA family, shown in the original skipping-rope representation and based on the previously reported consensus model^1^. Circle colors indicate nucleotide occupancy at the corresponding positions in the analyzed RF02924 alignment: red, ≥97%; black, ≥90%; gray, ≥75%; and open, ≥50%. The colors of nucleotide symbols indicate conservation of the specified nucleotide or nucleotide class: red, ≥97%; black, ≥90%; and gray, ≥75%. Base pairs exhibiting statistically significant covariation identified by R-scape are highlighted in green. R and Y denote purine and pyrimidine, respectively. (**B**) Secondary-structure diagram and residue numbering of the engineered two-strand env-2897 construct used for crystallization. Structural elements are colored as follows: P1, gray; P2 and the D-loop-like region, cyan; P3, green; P4 and the T-loop-like region, orange; and junctional nucleotides, magenta. The single 5-bromocytidine introduced for experimental phasing is indicated by a red box. The 5′ and 3′ termini of both RNA strands are indicated. (**C**) Parallel-eye stereoview of the env-2897 crystal structure (PDB: 9G4P), colored as in (**D**). The experimental electron-density map obtained after Br-SAD phasing and density modification is contoured at 1.2 σ. The final model contains 66 of the 68 nucleotides in the crystallization construct; C17 and G18 were not modeled because they lacked interpretable electron density.

A metagenome-derived representative, env-2897, was selected for crystallographic analysis (Figure S1B). A two-strand construct incorporating these elements and a single 5-bromocytidine substitution enabled structure determination at 2.9 Å resolution by bromine single-wavelength anomalous dispersion phasing (Br-SAD) (Figures 1B). The final model contains 66 of the 68 nucleotides; C17 and G18 were not modeled because they lacked interpretable electron density (Figure 1C; Figures S1D–S1F; Table S2).

Despite its unusual secondary-structure connectivity, env-2897 adopts a compact L-shaped fold resembling canonical tRNA (Figure 1C). Coaxial stacking of P1 and P4 generates an acceptor/T-arm-like segment, while P2 and P3 form a second arm occupying the position of the tRNA D/anticodon arm (Figures 1B and 1C). Contacts between the D-loop-like and T-loop-like regions organize the elbow and define the relative orientation of the two arms. The predicted long-range pseudoknot forms part of this interloop interface, directly linking the conserved secondary-structure elements to the tRNA-like fold. On the basis of this architecture, the skipping-rope family is designated tRNA-like RNA 1 (tLRNA-1).

Sequence conservation is concentrated around the four-way junction and elbow, where conserved nucleotides participate in an interconnected network of base-pairing and stacking interactions (Figures S1C-S1F; Table S2). By comparison, the P3 region, which occupies the position of the anticodon arm, displays substantially greater sequence variation(Figure S1A). This distribution is consistent with stronger evolutionary constraints on the tertiary core than on peripheral elements and identifies the junction and elbow as the principal sites for comparison with canonical tRNA.

### tLRNA-1 forms a tRNA-like elbow through alternative tertiary interactions

In canonical tRNAs, tertiary contacts between the D and T loops establish the elbow and organize the L-shaped fold^10^. Comparison of env-2897 with Saccharomyces cerevisiae tRNA^Phe^ shows similar overall dimensions despite substantial differences in the interactions supporting this arrangement (Figures 2A and 2B). The acceptor/T-arm-like segment of env-2897 closely superimposes with its tRNA counterpart, with axial dimensions of 47.9 and 49.8 Å, respectively (Figure 2C).

**Figure 2.**
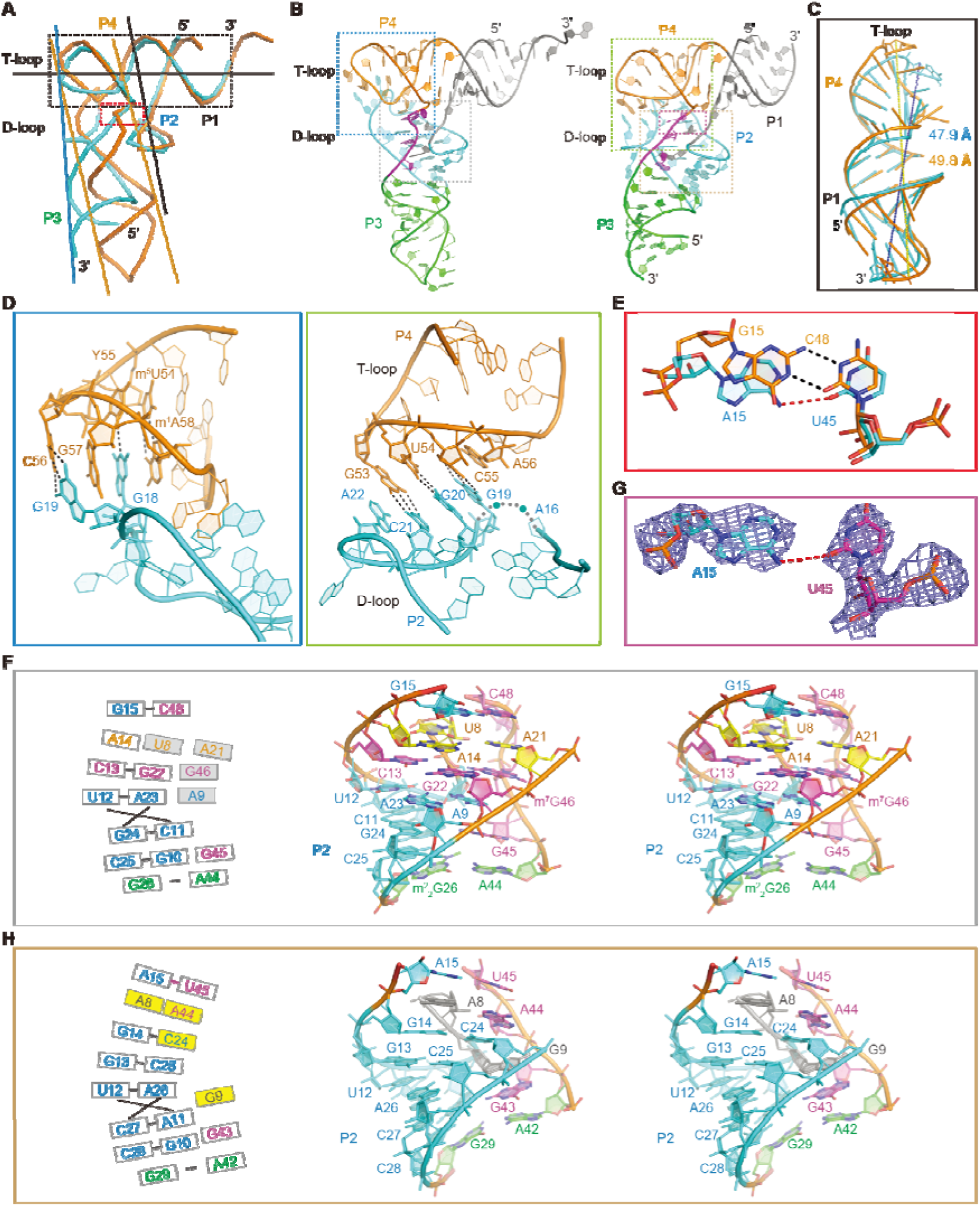
Structural comparison of canonical tRNA and the skipping-rope RNA. (**A**) Superposition of the env-2897 skipping-rope RNA structure (cyan; PDB: 9G4P) and *Saccharomyces cerevisiae* tRNA^Phe^ (orange; PDB: 1EHZ). The alignment was performed using conserved backbone atoms from selected elbow-associated regions, yielding an RMSD of 1.374 Å over 56 atoms. The corresponding regions are highlighted to illustrate the similarity in the overall L-shaped organization. Dashed boxes indicate the regions shown in (C) and (E). (**B**) Overall structures of env-2897 and *S. cerevisiae* tRNA^Phe^ shown side by side. Dashed boxes indicate the regions enlarged in (D) and (F-H). (**C**) Close-up comparison of the upper arm formed by P1/P4 in env-2897 and the acceptor/T-stem regions of canonical tRNA^Phe^. The coaxial stacking arrangement and corresponding helical dimensions are indicated. (**D**) Comparison of elbow-forming interactions between env-2897 (green box) and canonical tRNA^Phe^ (blue box). In env-2897, residues C17 and G18 lack interpretable electron density and are indicated by cyan spheres. A15 and A16 are omitted for clarity. Conserved D-loop/T-loop interactions in tRNA^Phe^, including the G18-Y55 and G19-C56 interactions, are shown for comparison. (**E**) Structural comparison of the Levitt pair-equivalent region. The A15–U45 interaction in env-2897 (cyan) is superimposed with the canonical reverse Watson–Crick G15–C48 Levitt pair in tRNA^Phe^ (orange). (**F**) Structural comparison of the junctional interaction networks in env-2897. The schematic representation (left) and stereoscopic structural view (right) highlight conserved residues and tertiary contacts involved in stabilizing the central junction. Residues are colored according to the structural elements shown in Figure 1B. (**G**) Close-up view of the A15-U45 interaction in env-2897. The electron-density map is shown as a gray mesh and contoured at 1.3 σ. Potential hydrogen-bonding interactions are indicated by dashed lines. (**H**) Structural comparison of the corresponding junctional region in canonical tRNA^Phe^. The schematic representation (left) and stereoscopic structural view (right) highlight the canonical tertiary interactions, including base triples and stacking interactions, that stabilize the tRNA elbow.

The similar elbow geometry is supported by different molecular contacts. In tRNA^Phe^, the T-loop contains the conserved U54–A58 reverse Hoogsteen pair, and the G18-Ψ55 and G19-C56 interactions connect the D and T loops. Together with the continuous stacking of A58, G18, G57, and G19, these contacts organize the canonical elbow^10^ (Figure 2D). The corresponding interface in tLRNA-1 instead contains three cis Watson-Crick/Watson-Crick pairs: G53-C21, U54-G20, and C55-G19 (Figure 2D). These contacts bring the two loop-like regions into a tRNA-like arrangement without reproducing the canonical D/T-loop interaction network.

A similar distinction occurs at the position of the Levitt pair. Canonical tRNAs contain a conserved G15-C48 reverse Watson-Crick pair within the junctional core^11^ (Figure 2E). In tLRNA-1, A15 and U45 occupy comparable spatial positions but do not reproduce this pairing geometry. The closest potential hydrogen-bonding contact, between A15(N6) and U45(O2), measures approximately 3.6 Å and provides limited support for a stable base-pairing interaction (Figures 2E and 2G). The local junction geometry is therefore maintained without an equivalent Levitt pair.

Around the D-stem of tRNA^Phe^, the U12-A23-A9, C13-G22-m^7^G46, and A14-U8-A21 base triples connect the helix to the tertiary core, together with stacking interactions extending from the Levitt pair through the D-stem (Figure 2H). These triples are absent from tLRNA-1. Instead, P2 contains the Watson-Crick pairs U12-A26, G13-C25, and G14-C24, with conserved A8 and A44 positioned on opposite sides of the helix to extend stacking through the junction (Figure 2F). The G29-A42 interaction further organizes the lower end of the core. Collectively, these contacts from tLRNA-1 maintain a tRNA-like elbow and junction without the canonical D/T-loop contacts, Levitt pair, or D-stem base-triple network.

### RCd1 accumulates as a highly stable processed RNA in *C. difficile*

RCd1 is a native member of the skipping-rope family in the human pathogen *Clostridioides difficile*. Originally identified by genome-wide transcriptional start site mapping, it is an abundant intergenic RNA located between CD630_1894 and CD630_1895, two genes of unknown function^7^. Previous work showed that Hfq depletion affects RCd1 accumulation and demonstrated RCd1-Hfq association by electrophoretic mobility shift assays *in vitro* and RNA co-immunoprecipitation followed by PCR analysis *in vivo*^8,9^. RCd1 is conserved among sequenced *C. difficile* strains. Northern blotting and RNA-seq detected abundant expression in both the laboratory strain 630Δerm and the epidemic ribotype 027 strain R20291 across multiple growth phases (Figures 3A and 3B), consistent with published transcriptomic data from additional strains and growth conditions^12^.

**Figure 3.**
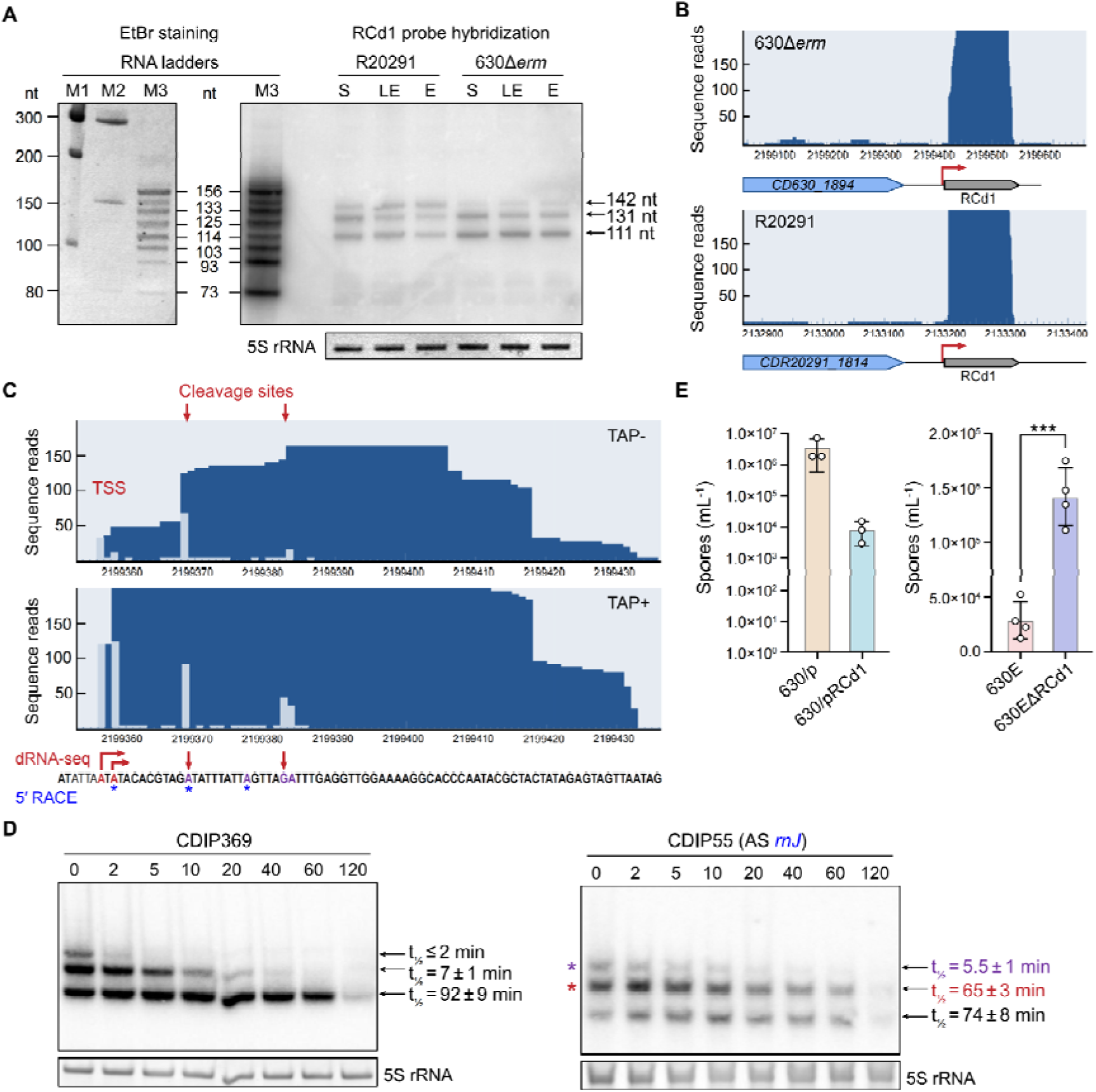
A naturally expressed skipping-rope RNA undergoes distinct processing and influences sporulation in *C. difficile*. (**A**) Northern blot detection of RCd1 transcripts in C. difficile strains 630Δerm and R20291 during exponential (E), late exponential (LE), and stationary (S) growth phases. Transcript sizes were estimated using in vitro-transcribed RNA markers. 5S rRNA served as a loading control. (**B**) RNA-seq read mapping of the RCd1 locus in 630Δerm and R20291 strains. RCd1 is located in an intergenic region downstream of CD630_1894/CDR20291_1814. Red arrows indicate the mapped transcription start site. (**C**) Mapping of RCd1 transcript boundaries and processing sites using differential RNA-seq and 5′ RACE. The primary transcription start site is indicated by the red arrow, and candidate processing sites are indicated by red arrows. Blue asterisks denote 5′ RACE products. (**D**) RCd1 transcript stability and RNase-dependent processing. RNA decay was measured after rifampicin treatment in control strain CDIP369 and RNase J-depleted strain CDIP55. Half-lives of the 142-, 131-, and 111-nt RCd1 species were quantified from Northern blot analyses. 5S rRNA served as loading control. (**E**) Sporulation phenotypes associated with altered RCd1 levels. Left: spore formation efficiency following RCd1 overexpression. Right: early sporulation efficiency in the RCd1 deletion strain. Data represent mean ± SD from independent biological replicates.

Transcript boundaries were mapped by differential RNA-seq and 5′/3′ rapid amplification of cDNA ends (RACE). The RCd1 locus lies downstream of CD630_1894 in 630Δerm and CDR20291_1814 in R20291 (Figure 3B). The primary 5′ end coincides with the major transcription start site downstream of a σA-dependent promoter, whereas the 3′ end maps to the poly(U) tract of an intrinsic transcription terminator (Figure 3C; Fig S3). These boundaries support independent transcription of RCd1 rather than its production as part of the downstream transcript. Comparison of TAP− and TAP+ profiles, together with internal ends detected by RACE, further supports post-transcriptional processing and identifies candidate cleavage sites (Figure 3C; Fig S4).

Northern blotting detected three major RCd1 species of approximately 142, 131, and 111 nt (Figure 3A). Their sizes were estimated using commercial markers together with defined RNA standards of 73, 93, 103, 114, 125, 133, and 156 nt generated by T7 *in vitro* transcription. The standards were detectable by both ethidium bromide staining and probe hybridization, allowing the RCd1 bands to be assigned against the same reference transcripts (Figure 3A; Fig S3).

The three RCd1 species differed markedly in stability following rifampicin-mediated transcriptional arrest. The ~ 142-nt species decayed rapidly, with a half-life of ≤2 min, whereas the ~131-nt species had a half-life of 7 ± 1 min. In contrast, the predominant ~111-nt species persisted with a half-life of 92 ± 9 min (Figure 3D). To assess the contribution of RNA-processing and decay factors, RCd1 turnover was measured following depletion of Hfq, RNase III, RNase J, or RNase Y^13^ (Figure S3).

Hfq or RNase III depletion had comparatively little effect on the RCd1 decay profile (Figure S3). RNase J depletion, however, altered the relative abundance of the RCd1 species and increased the half-life of the ~131-nt intermediate from 7 ± 1 to 65 ± 3 min; the half-life of the ~142-nt species increased to 5.5 ± 1 min (Figure 3D). RNase Y depletion produced a weaker effect, extending the half-life of the intermediate species to approximately 17 min and that of the longest species to approximately 5 min (Figure S3B; Fig. SY). The predominant ~111-nt RNA remained highly stable under all tested conditions. Candidate processing sites are mapped onto a predicted RCd1 structure in Fig. S3. These results implicate RNase J, and to a lesser extent RNase Y, in the turnover of longer RCd1 species but do not identify the nucleases that directly generate the stable short form.

All three RCd1 species are longer than typical mature tRNAs, which are approximately 75 nt and commonly undergo 5′ and 3′ maturation by RNase P and RNase Z, respectively^14^. RCd1 thus combines a tRNA-like structural assignment with a distinct transcript-size and turnover profile, accumulating predominantly as a highly stable processed RNA. Whether canonical tRNA-processing enzymes also contribute to its maturation remains undetermined.

### RCd1 negatively regulates sporulation in *C. difficile*

Previous work identified RCd1 as an Hfq-associated RNA implicated in sporulation control^9^. Its contribution to this process was assessed using an RCd1 deletion strain (630EΔRCd1) and a strain carrying an inducible RCd1 expression plasmid (630/pRCd1). After 24 h in sporulation medium, RCd1 overexpression reduced spore formation by approximately 1,000-fold relative to the empty-vector control, consistent with the previously reported inhibitory phenotype^9^ (Figure 3E). Conversely, the deletion strain produced approximately 10-fold more spores than the corresponding control strain after 7.5 h (Figure 3F). These opposing effects of RCd1 overexpression and deletion support a role for RCd1 in limiting sporulation under the conditions tested.

### DUF2693-FD and salivarius-1 RNAs share a tRNA-like core architecture

The arrangement of helices in tLRNA-1 provided a structural signature for searching other bacterial RNA families: an acceptor-stem-like helix directly adjacent to a T-loop-like stem-loop. A survey of structured RNAs in Rfam using this signature identified DUF2693-FD (RF02926) and salivarius-1 (RF03030) as candidate tRNA mimics, designated tLRNA-2 and tLRNA-3, respectively^2^ (Figures 4A-4C). Crystal structures of representatives from *Bacteroides dorei* and *Lactobacillus salivarius* were determined at 2.5 Å and 2.0 Å resolution, respectively. Both structures reproduce the upper-half architecture and elbow of canonical tRNA despite differences in sequence and secondary-structure topology (Figures 4D-4F).

**Figure 4.**
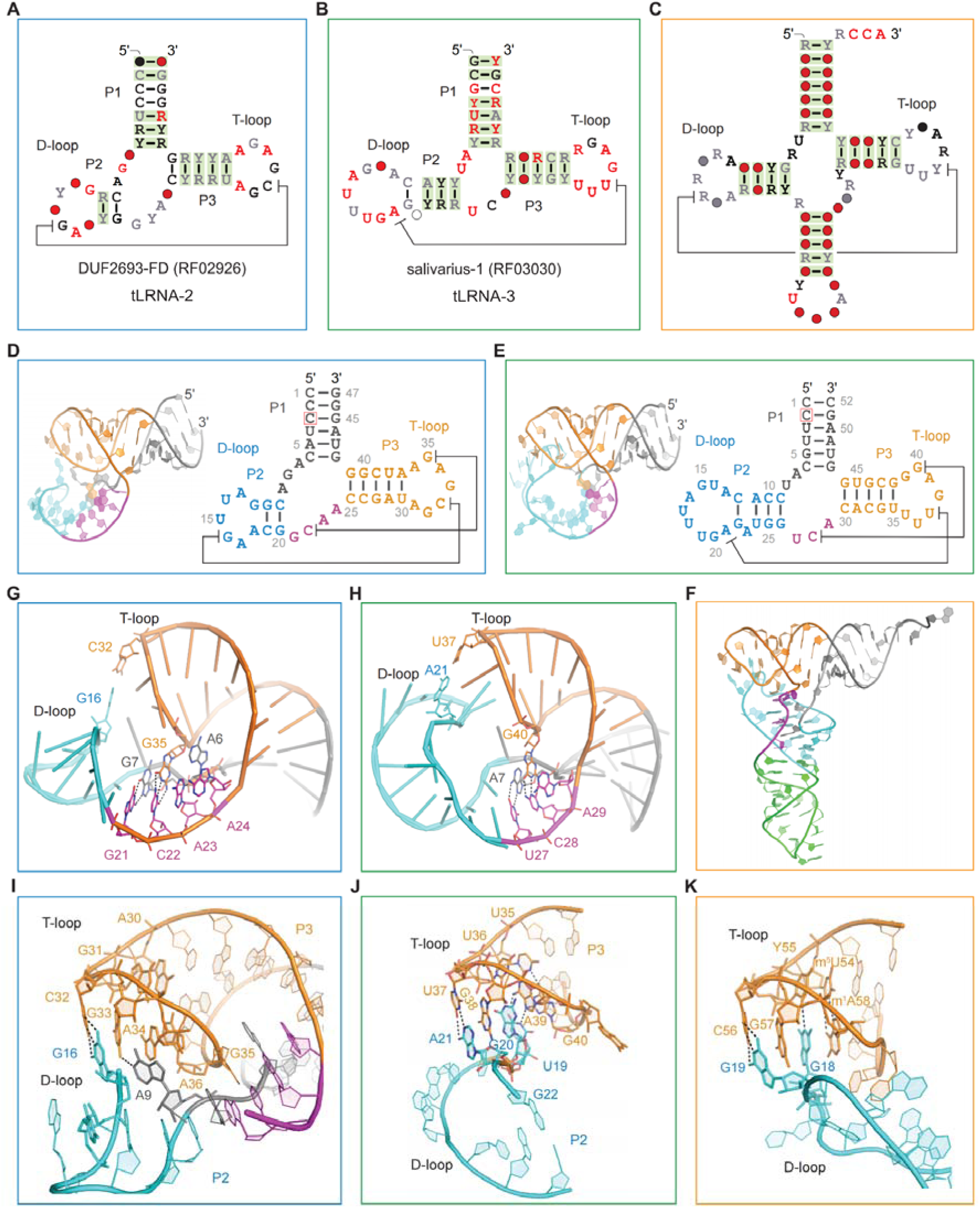
Independent bacterial RNA families converge on a conserved tRNA-like architecture and elbow organization. (**A-C**) Secondary-structure representations of representative bacterial tRNA-like RNA (tLRNA) families and canonical tRNA. (**A**) DUF2693-FD RNA family. (**B**) Salivarius-1 RNA family. (**C**) Canonical *Saccharomyces cerevisiae* tRNA^Phe^. Structural regions corresponding to the acceptor arm, D-loop-like region, and T-loop-like region are indicated. (**D-F**) Overall structures and structural organization of bacterial tLRNAs. (**D**) Crystal structure and secondary-structure representation of tLRNA-2. (**E**) Crystal structure and secondary-structure representation of tLRNA-3. (**F**) Structural comparison of tLRNA-2, tLRNA-3, and canonical tRNA^Phe^ (PDB: 1EHZ), highlighting the conserved spatial organization of the tRNA-like architecture despite distinct RNA topologies. (**G-H**) Close-up views of the central junction regions of tLRNA-2 and tLRNA-3. These regions illustrate the distinct tertiary interaction networks used by each RNA family to stabilize their tRNA-like folds. (**I-J**) Comparison of the elbow regions of tLRNA-2 and tLRNA-3. Although these RNAs lack the canonical tRNA D-loop/T-loop interaction network, both establish compact elbow-like structures through family-specific tertiary contacts. (**K**) Canonical D-loop/T-loop elbow of *S. cerevisiae* tRNA^Phe^ (PDB: 1EHZ), shown for comparison. Conserved tertiary interactions involving the D- and T-loop regions define the classical tRNA elbow architecture.

In tLRNA-2, coaxially stacked P1 and P3 form the acceptor/T-arm-like segment, with P2 projecting from this axis as a D-arm-like element (Figure 4D). The junction contains a G7-G21 cis Watson-Crick/Hoogsteen pair and a C22-G35 cis Watson-Crick/Watson-Crick pair within a network of noncanonical contacts (Figure 4G). Interactions between the P2 and P3 loops complete the elbow: C32 contacts G16, while hydrogen bonds involving G33-A9 and A36-A30 further organize the interface (Figure 4I). These contacts position the two loops in a tRNA-like arrangement through a local interaction network distinct from that of canonical tRNA.

A comparable helical arrangement is present in tLRNA-3, where stacked P1 and P3 form the acceptor/T-arm-like segment and P2 approaches the P3 loop to form the elbow (Figure 4E). Its junction is organized by A29 stacking on the C28-G40 pair and by a U27-A7 cis Watson-Crick/Hoogsteen interaction (Figure 4H). At the interloop interface, U37 pairs with A21 through Watson-Crick contacts, while G20 intercalates between G38 and A39 and forms a trans Watson-Crick/Sugar-edge interaction with U36 (Figure 4J). The resulting stacking network spans the P2–P3 interface and reproduces the spatial organization of the canonical D/T-loop elbow using different nucleotide identities and pairing geometries (Figures 4J and 4K).

Across the three families, an acceptor/T-arm-like helical scaffold is associated with a compact elbow assembled through family-specific base-pairing and stacking interactions. The shared geometry therefore extends beyond skipping-rope RNA, identifying a common structural organization among bacterial ncRNAs with distinct secondary structures.

### A shared architectural core guides the identification of candidate tRNA-like RNAs

Structural comparison localizes the common core of tLRNA-1, tLRNA-2, and tLRNA-3 to the upper half of the tRNA fold, encompassing the acceptor/T-arm-like scaffold and elbow. In canonical tRNA, this region provides recognition surfaces for multiple tRNA-binding and processing proteins^15,16^, including structural landmarks used by RNase P and RNase Z to position cleavage at the 5′ and 3′ ends, respectively^16–18^. Comparison with previously characterized tRNA-like RNAs further supports this region as a recurrent component of tRNA mimicry^3,6,19–22^.

The structural comparison suggested three features for a more explicit search signature: an acceptor-stem-like helix of approximately 5-8 bp, most commonly 7 bp; an adjacent T-stem-loop-like element, typically containing a 5-bp stem and a 7-nt loop; and direct connectivity between these elements without intervening nucleotides (Figures 5A-5D). A UUYRANY-like sequence pattern frequently occurs in the T-loop-like region (R, purine; Y, pyrimidine), providing an additional feature for candidate assessment. Application of these criteria to the 4,178 RNA families in Rfam 15.0 identified further candidate tLRNAs with the same core organization^2^.

**Figure 5.**
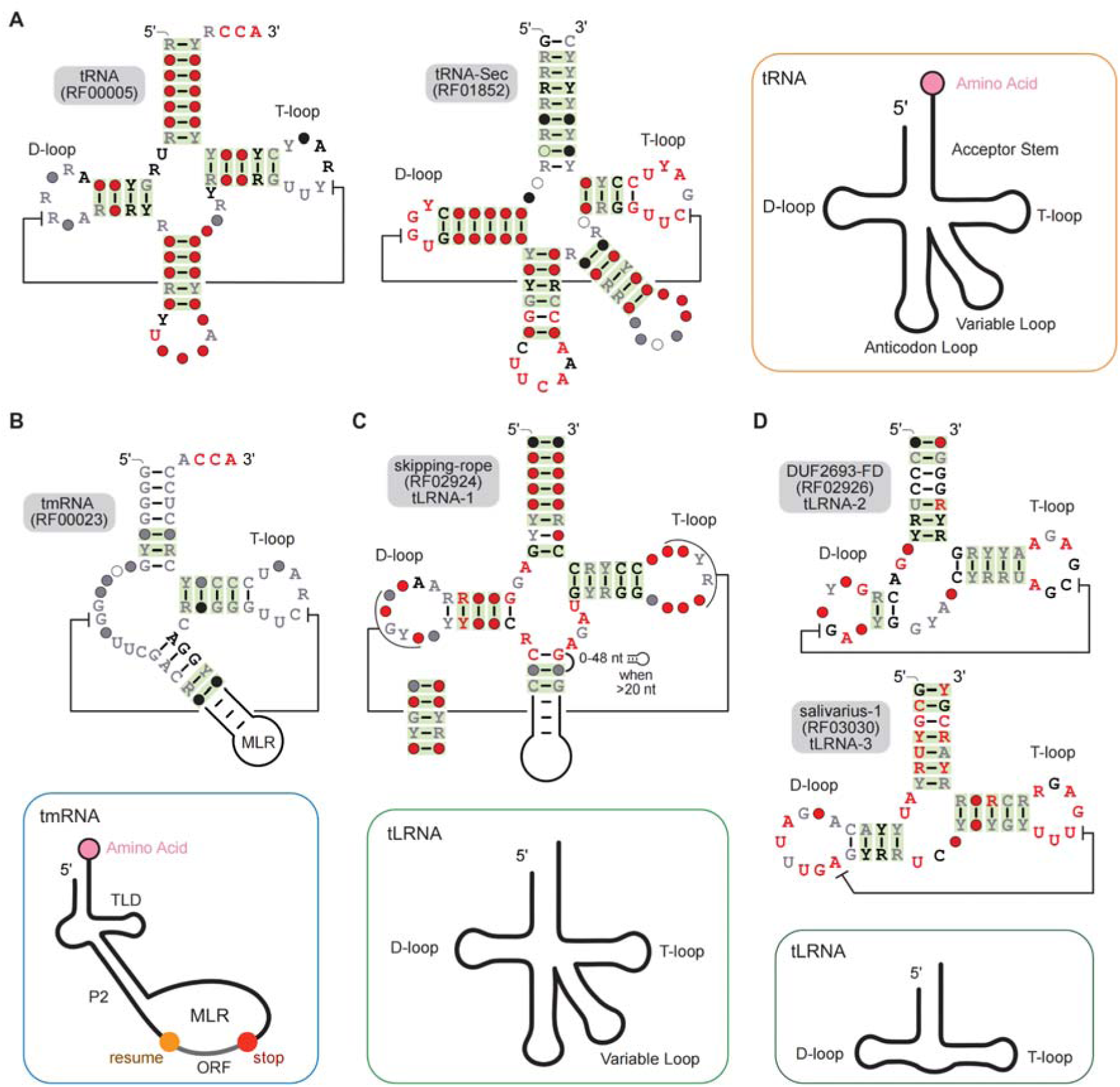
Topological evolution and secondary structural blueprints of the tRNA-like RNA (tLRNA) family compared to functional tRNAs. (**A**) Secondary structure blueprints of canonical tRNA (RF00005) and tRNA-Sec (RF01852). The inset (right) illustrates the classical cloverleaf topology, highlighting the spatial relationship between the upper half (Acceptor Stem and T-loop) and the decoding lower half (Anticodon loop). Highly conserved residues crucial for elbow formation are highlighted in red (T-loop) and dark grey (D-loop). (**B**) Secondary structure of bacterial tmRNA (RF00023). While adopting a tRNA-like domain (TLD) to interact with translation factors (e.g., EF-Tu) and RNases, the lower half is entirely replaced by an mRNA-like region (MLR) dedicated to ribosome rescue (inset). (**C**) Secondary structure of the skipping-rope RNA (RF02924; tLRNA-1). Representing a transitional architectural state, tLRNA-1 preserves the canonical upper half and core motif but entirely lacks the anticodon stem-loop, replacing it with a variable non-coding structural extension to serve as a post-transcriptional regulator. (**D**) Secondary structure blueprints of the highly minimalized DUF2693-FD and salivarius-1 motifs. The inset (bottom right) depicts the defining tLRNA topological paradigm: the extreme truncation of the canonical tRNA fold to its bare essential “core motif.” By abandoning the anticodon domain and strictly retaining the Acceptor Stem and T-loop equivalents, these RNAs autonomously form the highly stable, recumbent L-shaped “elbow” strictly optimized for protein binding rather than translation.

This core is embedded within RNAs of substantially different complexity (Figures 5A–5D). In canonical tRNAs, it is integrated with the D arm, anticodon arm, and variable region, whereas in tmRNA the tRNA-like domain forms part of a larger molecule containing additional structural elements and an mRNA-like coding region (Figures 5A and 5B). tLRNA-1 connects the corresponding scaffold to an alternative lower-half architecture comprising the P2/P3 arm and variable region (Figure 5C). By comparison, tLRNA-2 and tLRNA-3 are more compact, consisting predominantly of the acceptor/T-arm-like segment and the elements forming the elbow (Figure 5D). These comparisons distinguish the shared core from variable peripheral regions and provide a structural basis for identifying candidate tRNA mimics that lack the complete canonical cloverleaf.

## DISCUSSION

The identification of three bacterial RNA families as tRNA mimics reveals a shared architectural core that is not readily apparent from their secondary structures. Across these families, different patterns of base pairing and stacking support a similar tRNA-like elbow and helical scaffold. This finding separates conservation of three-dimensional geometry from conservation of the interactions that produce it. It also extends the basis for recognizing tRNA mimicry beyond resemblance to the canonical cloverleaf: RNAs with divergent sequences and connectivity can retain a common structural organization.

The shared core provides a practical link between experimentally determined structures and the identification of additional RNA families. Features extracted from the skipping-rope structure guided the recognition of further tRNA mimics, demonstrating how one structure can inform the classification of otherwise uncharacterized ncRNAs. The corresponding region of canonical tRNA participates in recognition by multiple proteins, including processing enzymes^16,17^. Its occurrence in other RNAs therefore suggests potential access to tRNA-recognition pathways. However, a shared fold does not establish common binding partners or biological activities. The core identified here provides a basis for generating testable functional hypotheses rather than assigning a single function to all tLRNAs.

RCd1 places this structural framework in the context of a native regulatory RNA. Its high stability and inhibitory effect on sporulation raise the possibility that a tRNA-like scaffold contributes to the persistence or molecular interactions of a bacterial regulator. These properties could reflect protection by the folded RNA itself, association with cellular proteins, or both. The established interaction of RCd1 with Hfq also leaves open a role for sequence-directed recognition of target transcripts^9^. Whether its tRNA-like core contributes directly to stability or sporulation control remains to be determined, as do the enzymes responsible for producing the stable processed species. Disrupting the core while monitoring RNA abundance, partner binding, and regulatory activity should help distinguish these possibilities.

More broadly, this work illustrates why experimental RNA structures remain essential for interpreting the expanding repertoire of ncRNA sequences. Protein annotation benefits from extensive catalogs of structurally and functionally characterized domains, which provide a foundation for inference from sequence. Comparable relationships remain much less developed for many structured ncRNA families. Building this foundation requires experimentally anchored connections among sequence, secondary structure, three-dimensional architecture, and biological activity. The present study contributes one such connection by translating an unexpected fold into recognizable features that identify further structural relatives. As these relationships accumulate across diverse RNAs, sequence and secondary structure should become increasingly informative about potential molecular partners and functions. Structural characterization thus has value beyond the individual RNA: it establishes the reference framework needed to turn newly discovered sequences into biologically interpretable hypotheses.

## Limitations of the study

Several limitations should be considered when interpreting our findings. First, the structures of tLRNA-1, tLRNA-2, and tLRNA-3 were determined using individual representatives of each RNA family under crystallization conditions, and the tLRNA-1 construct contained an engineered two-strand design and a 5-bromocytidine substitution for structure determination. Although the crystallographic structures are consistent with the conserved secondary-structure and covariation patterns of their respective RNA families, additional representatives and solution-state analyses will be required to establish the extent of conformational variation within each family and under physiological conditions. Second, although RCd1 is processed into multiple RNA species and depletion of RNase J and RNase Y alters the stability of the longer forms (Figure 3D and Figure S3), these experiments do not identify the nuclease or nucleases that directly generate the predominant ~ 11-nt species. Contributions from canonical tRNA-processing enzymes, including RNase P and RNase Z, therefore remain possible. Third, the reciprocal effects of RCd1 overexpression and deletion support a role for RCd1 in modulating sporulation in C. difficile (Figures 3E and 3F), but the direct molecular targets and mechanism linking RCd1 to the sporulation program remain unknown. In particular, whether RCd1 functions primarily through Hfq-dependent RNA–RNA interactions, through recognition of its tRNA-like architecture by cellular proteins, or through a combination of these mechanisms remains to be determined. Finally, biological functions have not yet been established for tLRNA-2, tLRNA-3, or the additional candidates identified by the core-motif search. The structural criteria used to identify candidate tLRNAs may also preferentially recover RNAs closely resembling the currently characterized core and may miss more divergent tRNA-like architectures. Functional and biochemical characterization of these candidates will therefore be necessary to determine how broadly the structural convergence described here corresponds to a shared biological strategy.

## METHOD DETAILS

### RNA synthesis

RNA oligonucleotides were synthesized using ***t***-BDMS phosphoramidite chemistry^23^ on an Applied Biosystems 394 DNA/RNA synthesizer, as described previously^24^. Oligonucleotides containing 5-bromocytidine (ChemGenes) were deprotected in a 25% ethanol/ammonia solution at 20°C for 36 h. All oligoribonucleotides were then redissolved in a mixture of 100 μL of anhydrous DMSO and 125 μL of triethylamine trihydrofluoride (Aldrich) and incubated with agitation at 65°C for 2.5 h in the dark to remove the ***t***-BDMS protecting groups. After cooling on ice for 10 min, the RNA was precipitated by adding 1 mL of butanol, washed twice with 70% ethanol, and dissolved in double-distilled water. The RNA was further purified by denaturing polyacrylamide gel electrophoresis in the presence of 7 M urea. The full-length RNA product was visualized by UV shadowing, and the corresponding band was excised. RNA was electroeluted using an Elutrap Electroelution System (GE Healthcare) into a buffer containing 45 mM Tris-borate (pH 8.5) and 5 mM EDTA at 200 V for 8 h at 4°C. The recovered RNA was precipitated with ethanol, washed once with 70% ethanol, and dissolved in double-distilled water.

### Chemicals and Reagents

Unless otherwise specified, all chemicals and reagents were purchased from Sigma-Aldrich at the highest purity available.

### Crystallization, structure determination, and refinement

RNA at a concentration of 0.5-0.8 mM in a buffer containing 5 mM HEPES (pH 7.6) and 100 mM KCl was heated to 95°C for 1 min and then slowly cooled to 20°C. MgCl_2_ was subsequently added to a final concentration of 2 mM. Crystallization conditions are listed in Table S1. X-ray diffraction data were collected at Diamond Light Source (Harwell, UK) on the beamlines specified in Table S1. Diffraction data were processed using xia2 version 0.5.270^25^. High-resolution cutoffs were selected based on CC_1/2_ statistics^26^ and inspection of the electron-density maps. All three structures reported in this study (PDB IDs: 9G4P, 9G4R, and 9G4Q) were determined by bromine single-wavelength anomalous diffraction (Br-SAD) phasing. Bromine sites were located using AutoSol in the PHENIX suite, and experimental electron-density maps were calculated following phasing and density modification^27^. Models were manually built and adjusted in Coot^28^, followed by iterative cycles of manual rebuilding in *Coot* and crystallographic refinement with phenix.refine^29^. Model geometry was assessed using MolProbity^30^ and the validation tools in Coot, and model fit to the electron density was evaluated in *Coot*.

### Sequence alignment and analysis

Sequences of skipping-rope, DUF2693-FD, and Salivarius-1 RNAs were retrieved from Rfam under accession numbers RF02924, RF02926, and RF03030, respectively. The sequence alignments were manually refined in Jalview using the crystal structure as a guide^31^. Nucleotide composition and covariation analyses were performed using Jalview.

### Plasmid and bacterial strain construction and growth conditions

C. difficile strains were grown in TY medium^32^ or brain heart infusion (BHI) medium (Difco) in an anaerobic chamber (Jacomex) under an atmosphere of 5% H, 5% CO_2_, and 90% N_2_. Where required, media were supplemented with cefoxitin (Cfx; 25 μg/mL) and thiamphenicol (Tm; 15 μg/mL). *E. coli* strains were grown in LB broth^33^ supplemented, where appropriate, with ampicillin (100 μg/mL) or chloramphenicol (15 μg/mL). Anhydrotetracycline (ATc) was used to induce the Ptet promoter in pRPF185 derivatives in *C. difficile*^34^. Unless otherwise specified, strains carrying pRPF185 derivatives were grown for 7.5 h in TY medium supplemented with 250 ng/mL ATc and 7.5 μg/mL Tm. Routine plasmid construction was performed using standard molecular cloning procedures. Deletion of the *RCd1* gene was performed using an allelic exchange approach based on a semi-suicidal plasmid carrying the *E. coli codA* gene, which encodes cytosine deaminase, as a counterselection marker^35^. The deletion plasmid was constructed by Gibson assembly for subsequent conjugation and homologous recombination in *C. difficile*. The resulting plasmid was introduced into *E. coli* HB101 (RP4), which was then conjugated with *C. difficile* 630Δerm. Transconjugants were selected by subculturing on BHI agar supplemented with Tm (15 μg/mL) and Cfx (25 μg/mL).

### RNA extraction, Northern blot and 5′/3′ RACE

Total RNA was extracted from C. difficile strains grown in TY medium at the exponential phase (4 h), late exponential phase (6 h), and onset of stationary phase (9.5 h), as previously described^36^. Northern blotting and 5′/3′ rapid amplification of cDNA ends (RACE) were performed as previously described^7^. To generate an RCd1-specific RNA size ladder, RNA fragments containing RCd1 sequences of different lengths were synthesized *in vitro* using T7 RNA polymerase and the MAXIscript kit (Thermo Fisher Scientific), with blunt-ended PCR products as templates, according to the manufacturer’s instructions. The length of the RCd1 transcript was estimated by comparison with both commercial and RCd1-specific RNA size ladders.

### Measurement of RNA decay by rifampicin assay

RNA decay was assessed using a rifampicin assay as previously described^13^. Briefly, to determine RCd1 RNA half-lives, *C. difficile* strains were grown at 37°C for 7.5 h in TY medium supplemented with 250 ng/mL ATc and 7.5 μg/mL Tm. Rifampicin was added to a final concentration of 200 μg/mL, and samples were collected at 0, 2, 5, 10, 20, 40, 60, and 120 min for RNA extraction and Northern blotting as described above.

### Sporulation assay

Overnight cultures grown in TY medium at 37°C were used to inoculate 20 mL of sporulation medium^37^. Where required, Tm (7.5 μg/mL) and ATc (250 ng/mL) were added for plasmid maintenance and induction of RCd1 expression from pRCd1, respectively. After 7.5 or 24 h of incubation, a 1-mL culture aliquot was collected and divided into two samples. To determine total colony-forming units (CFU), the first sample was serially diluted in liquid sporulation medium and plated on BHI agar supplemented with 0.1% taurocholate (Sigma-Aldrich) to promote spore germination^37^. To enumerate spores, the second sample was incubated at 65°C for 20 min to kill vegetative cells before plating on BHI agar supplemented with 0.1% taurocholate. The sporulation frequency was calculated as the ratio of spore CFU/mL to total CFU/mL.

## AUTHOR CONTRIBUTIONS

LH conceived the project. LH, OS and JW designed the experiments. LH and JW purified the RNAs, performed crystallographic experiments and bioinformatic analyses, and determined the structures. LH and OS performed RNA-seq analyses. J.W., L.H., D.M.J.L. and O.S. acquired funding. P.B. and O.S. performed all experiments related to RCd1. L.H., Y.X. and O.S. organized the figures and tables. J.W., L.H., P.B., O.S., Y.X., Y.H., Y.R., D.M.J.L., E.W. and Q.C.analyzed the data. L.H., O.S., JW, Y.X.,Y.H. and Y.R. contributed to writing the initial draft of the manuscript.

## SUPPLEMENTAL INFORMATION

Supplementary Figures S1-S4 and supplementary Tables S1-S2 are available.

## ACKNOWLEDGEMENTS

We thank Saira Ashraf for RNA synthesis. This work was supported by the Natural Science Foundation of Top Talent of SZTU (grant no. GDRC202418 to J.W.) and the Guangdong Science and Technology Department (2026B1212170033 and 2020B1212030004 to L.H.).This work was also supported by grants from Cancer Research UK (program grant A18604 to D.M.J.L.), from the Agence Nationale de la Recherche (ANR-11-IDEX-0003-02; “CloSTARn”, ANR-13-JSV3-0005-01 to O.S., “CdiffRib” ANR-22-CE15-0020-01 to O.S.; “CDefenseRNA”, ANR-24-CE35-2137-01 to O.S.; “Tartas”, ANR-24-CE15-2031-02 to O.S.). We thank Drs Yiliang Ding, Gillian Douce, Nicola Stanley-Wall and Helge Dorfmueller for valuable discussion. We thank Diamond for synchrotron time and the Wellcome Trust for the in-house diffractometer.

## Notes

### Competing Interest Statement

The authors have declared no competing interest.

