## Supplemental Information for "Structural analysis reveals three families of bacterial tRNA mimics"

### Three bacterial tRNA-like RNAs revealed by structural analysis

**Figure S1**

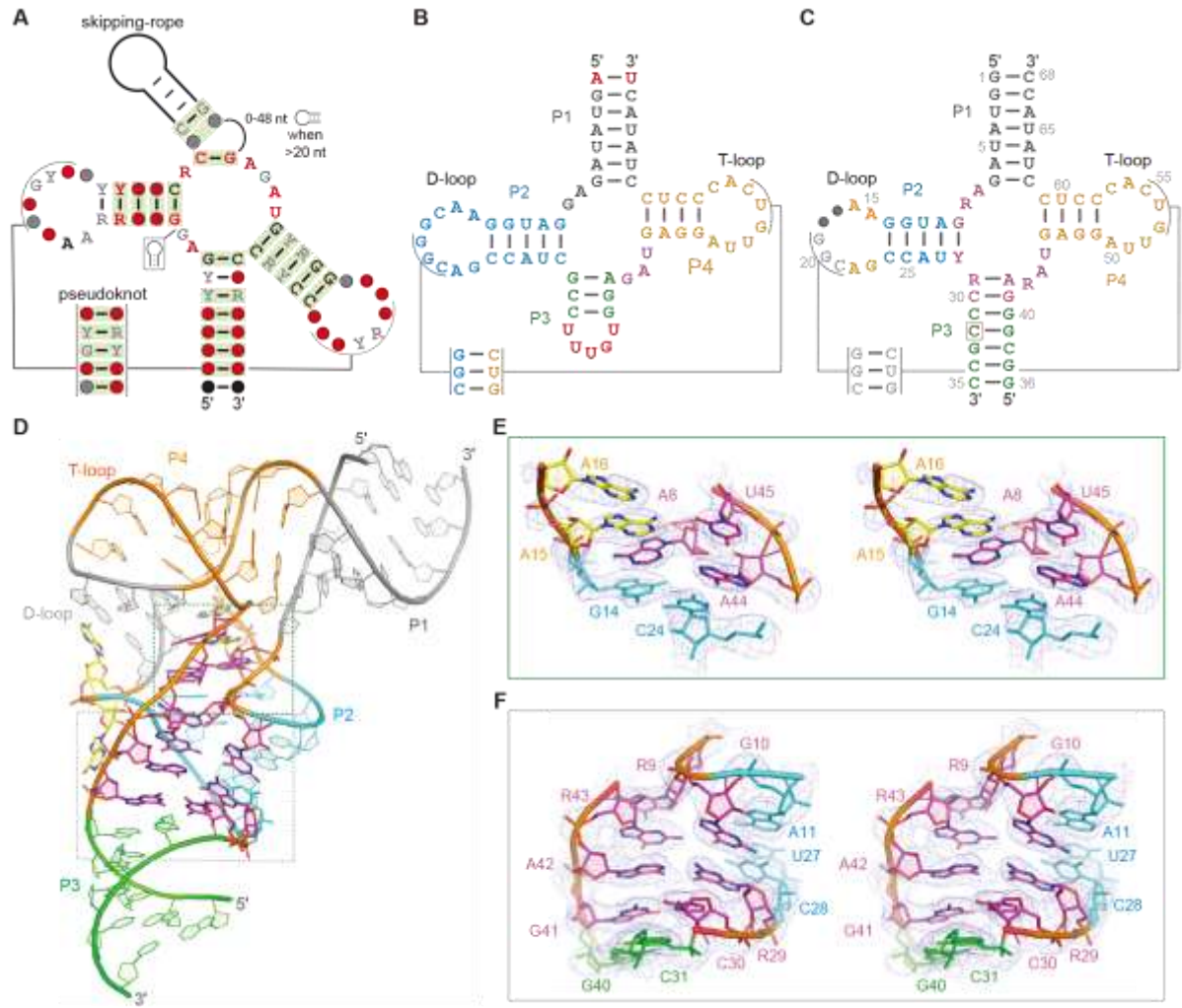

**Figure S1. Consensus models, construct design, and conserved structural features of the skipping-rope RNA.** (A) Consensus sequence and secondary-structure model of the skipping-rope RNA motif (Rfam RF02924). Base pairs with significant covariation identified by R-scape are highlighted in green. (B) Wild-type env-2897 skipping-rope RNA sequence. Nucleotides differing between the wild-type RNA and the crystallization construct are highlighted in red. (C) Conserved nucleotides surrounding the four-way junction. Positions with $\geq 97\%$ conservation in the analyzed RF02924 alignment are highlighted in magenta. (D) Overall crystal structure of the env-2897 skipping-rope RNA. Structural elements are colored as in Figure 1B. The regions shown in (E) and (F) are boxed. The density-modified Br-SAD experimental electron-density map is contoured at $1.2 \sigma$. Conserved nucleotides cluster around the four-way junction. (E) Close-up view of the conserved junctional residues A8, A44, and U45. (F) Close-up view of conserved residues involved in the junctional interaction network. Residue numbering corresponds to the

crystallized env-2897 construct

**Figure S2**

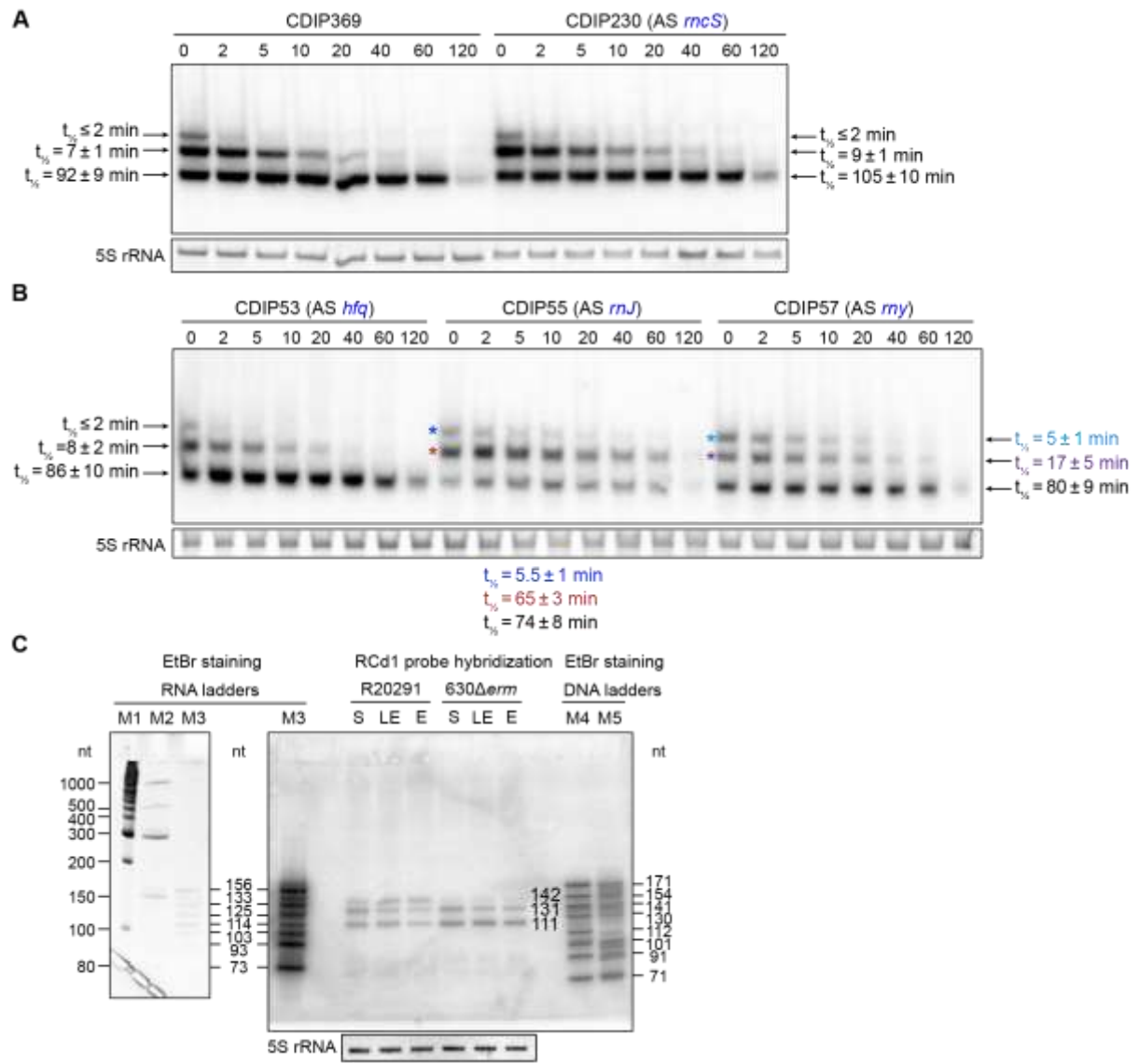

**Figure S2. Identification of ribonucleases involved in RCd1 transcript maturation, related to Figure 3.** (A) Effects of Hfq and RNase III depletion on RCd1 transcript stability. RCd1 decay kinetics were measured following rifampicin-mediated transcriptional arrest. Northern blot analysis was performed using an RCd1-specific probe, and transcript half-lives were calculated from decay profiles. 5S rRNA served as a loading control. (B) Effects of RNase J and RNase Y depletion on RCd1 transcript stability. RNase J-depleted (CDIP55) and RNase Y-depleted (CDIP57) strains were analyzed following rifampicin treatment. RNase J depletion resulted in strong stabilization of the 131-nt intermediate species, whereas RNase Y depletion produced a weaker stabilization effect on RCd1 precursor species. (C) Validation of RCd1 transcript species and size estimation using in vitro-transcribed RNA markers. The identified RCd1 transcript forms correspond to the major species detected by Northern blot analysis.

**Figure S3**

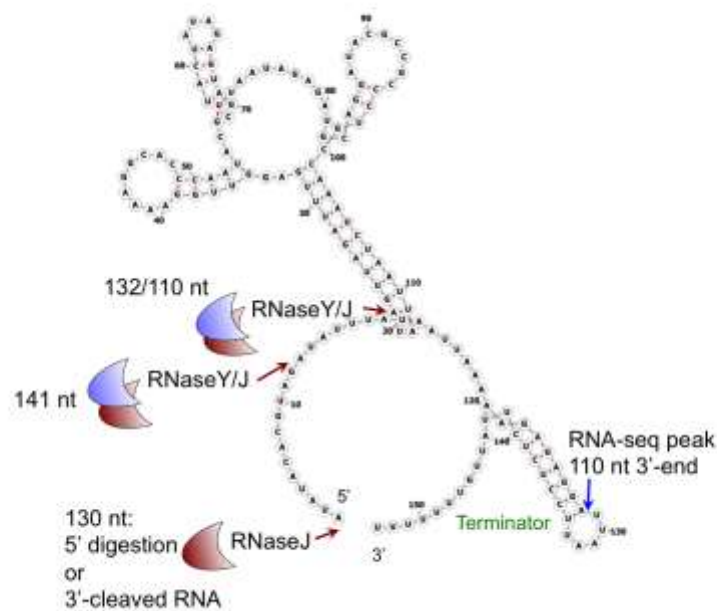

**Figure S3. RCd1 secondary structure prediction by RNAMfold software.** The positions of potential cleavage sites by RNase J with 5'-3' exoribonuclease and endoribonuclease activities and by endoribonuclease RNase Y are shown based on the RNA-seq, 5'/3' RACE and RNA stability analysis. The potential processing site in 3'-end was deduced from RNA-seq data. The 3'-end stem-loop structure corresponds to the rho-independent terminator.

**Figure S4**

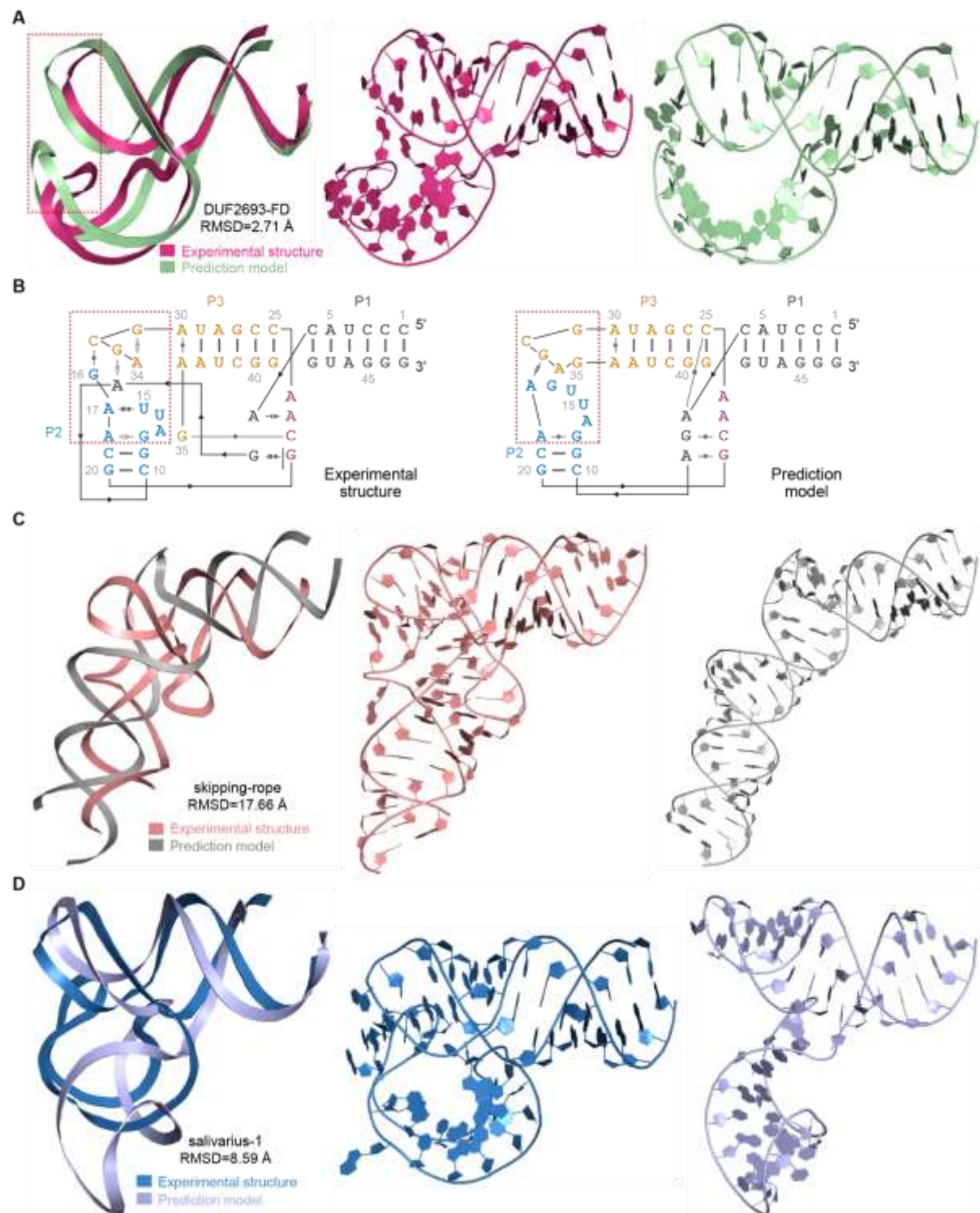

**Figure S4. Comparison of experimental structure and predicted structure by AlphaFold3.** (A) Superposition of the experimental (hot pink) and predicted structures (green) of DUF2693-FD, focusing on the overall conformation with an RMSD of 2.71 Å. The predictive model with the highest-ranking score is selected for comparative analysis and RMSD analysis is based on

all atoms. The red box highlights the D-T loop interaction region, which is also the location where the main differences exist between the two structures. **(B)** Crystal structure-derived secondary structure of DUF2693-FD: experimental (left) and predicted model (right). Noncanonical base pairs are indicated by Leontis–Westhof symbols<sup>1</sup>. Lines with embedded arrowheads denote 5'-to-3' chain connectivity. The red box zooms in on the detailed interactions in (A). **(C)** and **(D)** Overall structure superposition of experimental and predicted structures for skipping-rope (salmon/gray) and salivarius-1 (blue/light blue), revealing substantial structural discrepancies with RMSDs of 17.66 Å and 8.59 Å, respectively.

**Table S1. Crystallographic data collection and refinement statistics for the three bacterial tRNA-like RNAs.**

| RNA Type | Skipping-rope<br>(tLRNA-1) | DUF2693-FD<br>(tLRNA-2) | Salivarius-1<br>(tLRNA-3) |
| --- | --- | --- | --- |
| PDB ID | 9G4P | 9G4R | 9G4Q |
| Phasing | SAD | SAD | SAD |
| Data collection |  |  |  |
| Space group | $P 6_1$ | $P 6_5 2 2$ | $P 2_1 2_1 2_1$ |
| Cell dimensions |  |  |  |
| a, b, c (Å) | 113.5, 113.5, 59.3 | 110.8, 110.8, 74.8 | 49.4, 68.1, 99.3 |
| $\alpha, \beta, \gamma$ (°) | 90, 90, 120 | 90, 90, 120 | 90, 90, 90 |
| Wavelength | 0.9196 | 0.9159 | 0.9198 |
| Resolution (Å) | 98.28 – 2.90<br>(2.95 – 2.90) | 47.96 – 2.53<br>(2.57 – 2.53) | 56.17 – 1.99<br>(2.02 – 1.99) |
| Rmerge | 0.095 (1.091) | 0.058 (0.800) | 0.109 (3.50) |
| $\langle I \rangle / \langle \sigma(I) \rangle$ | 16.5 (2.3) | 17.7 (1.8) | 15.6 (1.4) |
| CC$_{1/2}$ | 1.00 (0.28) | 1.00 (0.69) | 1.00 (0.68) |
| Completeness (%) | 100.0 (100.0) | 100.0 (100.0) | 100.0 (98.5) |
| Redundancy | 19.7 (20.1) | 7.1 (7.3) | 25.3 (25.8) |
| Refinement |  |  |  |
| Resolution (Å) | 37.14 – 2.90<br>(3.00 – 2.90) | 47.96 – 2.53<br>(2.62 – 2.53) | 29.78 – 1.99<br>(2.06 – 1.99) |
| No. reflections | 9807 (977) | 9476 (923) | 23585 (2326) |
| Rwork / Rfree | 0.199 / 0.233 | 0.189 / 0.228 | 0.236 / 0.268 |
| No. atoms |  |  |  |
| Macromolecules | 1411 | 1027 | 2165 |
| Ligands | 4 | 32 | 48 |
| Solvent | N/A | 2 | 44 |
| Mean B-factors (Å<sup>2</sup>) |  |  |  |
| Macromolecules | 109.23 | 82.71 | 59.88 |
| Ligands | 116.34 | 88.08 | 63.67 |
| R.m.s. deviations |  |  |  |
| Bond lengths (Å) | 0.005 | 0.008 | 0.003 |
| Bond angles (°) | 0.82 | 1.25 | 0.60 |

Values in parentheses are for the highest-resolution shell.

**Table S2. Sequence conservation and nucleotide-pair composition of selected residues in the skipping-rope RNA family.**

| Base Pair<br>/Nucleotide | Pairing Type | Frequency (%) |
| --- | --- | --- |
| Watson-Crick pairs |  |  |
| G7-C62 | GC=92.8%, AU=5.6% | WC = 98.4 |
| G10-Y28 | GC=92.4%, GU=6.5% | GY=98.9% |
| C30-G41 | CG=97.7% | WC=98.9% |
| A15-U45 | AU=78.0%, GU=13.5%, CU=4.8%,<br>AG=1.9% | RU=91.5% |
| Non-canonical<br>/Single base |  |  |
| A8 | A=99.9% | A=99.9% |
| R9 | G=85.0%, A=13.7% | R=98.7% |
| R29-A42 | AA=50.2%, GA=48.9%, AG=0.3%, | RA=99.1% |
| R43 | G=82.7%, A=15.3%, U=1.8% | R=98.0% |
| A44 | A=99.4%, G=0.6% | R=100.0% |

Nucleotides colour magenta are >97% identical in all species analyzed around the four-way junction.
